# GPR183 regulates interferons and bacterial growth during *Mycobacterium tuberculosis* infection: interaction with type 2 diabetes and TB disease severity

**DOI:** 10.1101/2020.07.15.203398

**Authors:** Stacey Bartlett, Adrian Tandhyka Gemiarto, Minh Dao Ngo, Haressh Sajiir, Semira Hailu, Roma Sinha, Cheng Xiang Foo, Léanie Kleynhans, Happy Tshivhula, Tariq Webber, Helle Bielefeldt-Ohmann, Nicholas P. West, Andriette M. Hiemstra, Candice E. MacDonald, Liv von Voss Christensen, Larry S. Schlesinger, Gerhard Walzl, Mette Marie Rosenkilde, Thomas Mandrup-Poulsen, Katharina Ronacher

## Abstract

Oxidized cholesterols have emerged as important signaling molecules of immune function, but little is known about the role of these oxysterols during mycobacterial infections. We found that expression of the oxysterol-receptor GPR183 was reduced in blood from patients with tuberculosis (TB) and type 2 diabetes (T2D) compared to TB patients without T2D and was associated with TB disease severity on chest x-ray. GPR183 activation by 7α,25-hydroxycholesterol (7α,25-OHC) reduced growth of *Mycobacterium tuberculosis* (Mtb) and *Mycobacterium bovis* BCG in primary human monocytes, an effect abrogated by the GPR183 antagonist GSK682753. Growth inhibition was associated with reduced IFN-β and IL-10 expression and enhanced autophagy. Mice lacking GPR183 had significantly increased lung Mtb burden and dysregulated IFNs during early infection. Together, our data demonstrate that GPR183 is an important regulator of intracellular mycobacterial growth and interferons during mycobacterial infection.

**Graphical Abstract:** 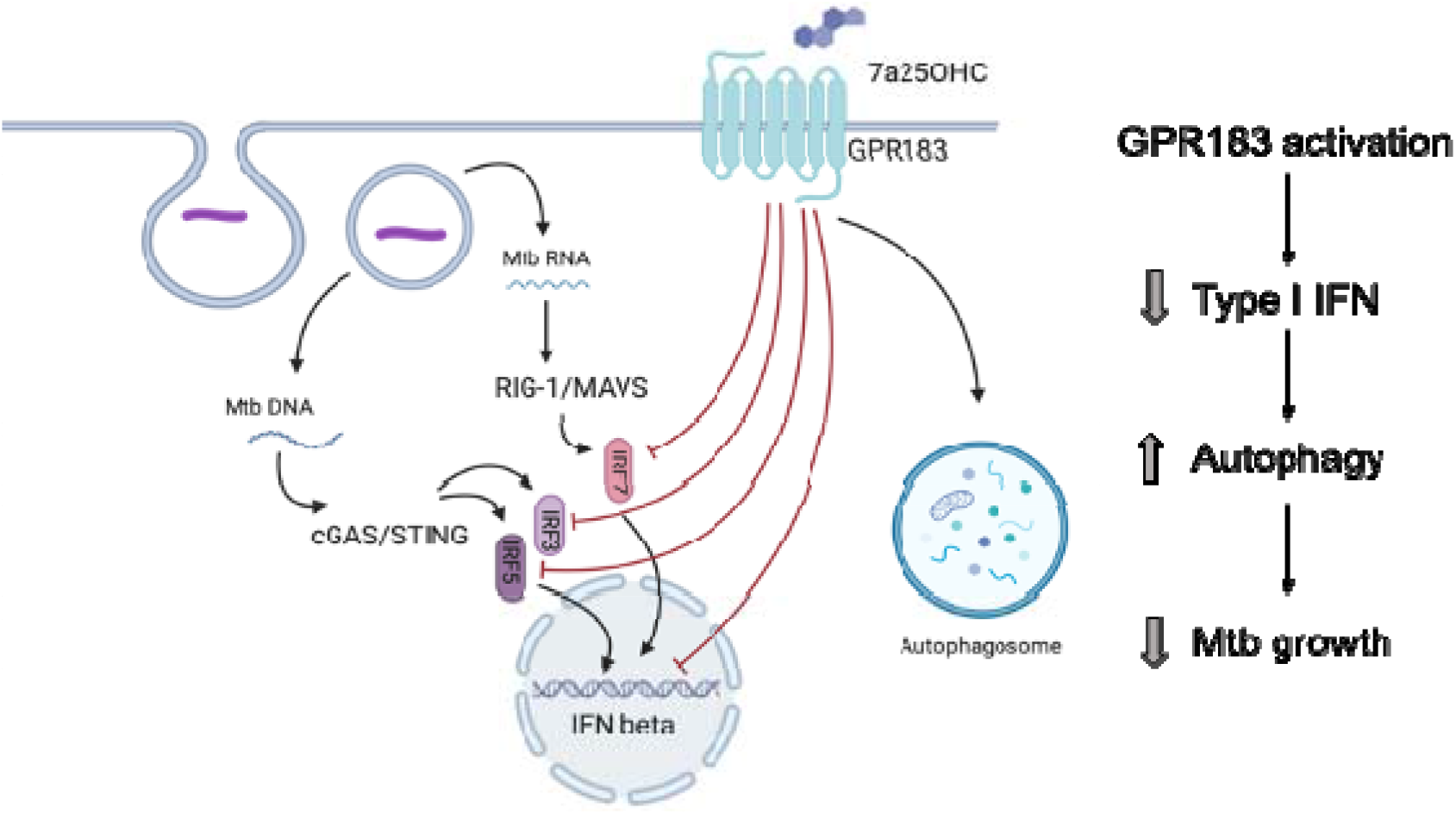

## Background

Patients with tuberculosis and type 2 diabetes (TB-T2D) co-morbidity have increased bacterial burden and more severe disease, characterized by higher sputum smear grading scores and greater lung involvement on chest x-ray compared to TB patients without T2D [1]. TB-T2D patients are also more likely to fail TB therapy and to relapse [1]. The reason for the increased disease severity has largely been attributed to hyperglycemia-mediated immune dysfunction, but hyperglycemia alone does not fully explain these observations [1, 2]. We recently showed that independent of hyperglycemia, cholesterol concentrations in T2D patients vary greatly across different ethnicities [3]. However, how cholesterol and its metabolites contribute to *Mycobacterium tuberculosis* (Mtb) infection outcomes remains to be elucidated.

To gain novel insights into the underlying immunological mechanisms of increased susceptibility of T2D patients to TB and to identify novel targets for host-directed therapy (HDT), we performed whole blood transcriptomic screens on TB patients with and without T2D and identified differential regulation of the transcript for oxidized cholesterol-sensing G protein-coupled receptor (GPCR), GPR183. Also known as Epstein Barr virus-induced gene 2 (EBI2), GPR183 is primarily expressed on cells of the innate and adaptive immune system [4–6]. Several oxysterols can bind to GPR183 with 7α,25-hydroxycholesterol (7α,25-OHC) being the most potent endogenous agonist [4, 7, 8]. GPR183 has been studied mainly in the context of viral infections [9], immune cells [4, 5, 7, 10–16], and astrocytes [17, 18]; and facilitates the chemotactic distribution of lymphocytes, dendritic cells and macrophages to secondary lymphoid organs [10, 13, 14, 19, 20]. Little is known about the biological role of GPR183 in the context of bacterial infections, including TB. We show here that GPR183 is a key regulator of intracellular bacterial growth and type-I IFN production during mycobacterial infection and reduced GPR183 expression is associated with increased TB disease severity.

## Methods

### Study participants

TB patients and their close contacts were recruited at TB clinics outside Cape Town (South Africa). TB diagnosis was made based on positive GeneXpert MTB/RIF (Cepheid; California, USA) and/or positive MGIT culture (BD BACTED MGIT 960 system, BD, New Jersey, USA) and abnormal chest x-ray. Chest x-rays were scored, based on Ralphs score [21], by two clinicians independently. Participants with LTBI were close contacts of TB patients, who tested positive on QuantiFERON-TB Gold in tube assay (Qiagen, Hilden, Germany). All study participants were screened for T2D based on HbA1c ≥ 6.5% and random plasma glucose ≥ 200 mg/dL or a previous history of T2D. Further details are available in the supplementary materials.

### RNA extractions and Nanostring Analysis

Total RNA was extracted from cell pellets collected in QuantiFERON-TB gold assay tubes without antigen using the Ribopure Ambion RNA isolation kit (Life Technologies, California, USA), and eluted RNA treated with DNase for 30 min. Samples with a concentration of ≥ 20 ng/μL and a 260/280 and 260/230 ratio of ≥ 1.7 were analyzed at NanoString Technologies in Seattle, Washington, USA. Differential expression of 594 genes, including 15 housekeeping genes, was performed using the nCounter GX Human Immunology kit V2. NanoString RCC data files were imported into the nSolver 3 software (nSolver Analysis software, v3.0) and gene expression was normalized to housekeeping genes.

### Cell culture

Peripheral blood mononuclear cells (PBMCs) were obtained from healthy donor blood by Ficoll-Paque (GE Healthcare, Illinois, USA) gradient centrifugation and monocytes (MNs) isolated using the Pan Monocyte Isolation kit (Miltenyi Biotec, Bergisch Gladbach, Germany), with >95% purity assessed by flow cytometry. MNs were plated onto Poly-D-lysine coated tissue culture plates (1.3 × 10^5^ cells/well) and rested overnight at 37°C/5%CO_2_ in RPMI-1640 medium supplemented with 10% heat-inactivated human AB serum (Sigma Aldrich, Missouri, USA), 2 mM L-glutamine and 1 mM sodium pyruvate before infection. THP-1 cells (ATCC #TIB-202) were differentiated with 25 ng/mL PMA for 48h and rested for 24h prior to infection.

### In vitro Mtb (H_7_R_v_)/M. bovis (BCG) infection

Mtb H_37_R_v_ or *M. bovis* BCG single cell suspensions were added at a multiplicity of infection (MOI) of 1 or 10 with/without 100 nM 7α,25-dihydroxycholesterol (Sigma Aldrich) and with/without 10 μM GSK682753 (Focus Bioscience, Queensland, Australia), followed by 2h incubation at 37°C/5%CO_2_ to allow for phagocytosis. Non-phagocytosed bacilli were removed by washing each well twice in warm RPMI-1640 containing 25 mM HEPES (Thermo Fisher Scientific). Infected cells were incubated (37°C/5%CO_2_) in medium with/without GPR183 agonist and/or antagonist and CFUs determined after 48h.

To quantify bacterial growth over time, CFUs at 48h were normalized to uptake at 2h. Percentages of mycobacterial growth were determined relative to untreated cells. For RNA extraction, MNs were lysed by adding 500 uL of TRIzol reagent. Further details are provided in the supplementary information.

### Western Blotting

THP-1 cells were infected with BCG with/without 100nM 7α,25-OHC and with/without 10 μM GSK682753 and lysed at 6 or 24h post infection (p.i.) in ice-cold RIPA buffer (150 mM sodium chloride, 1.0% Triton X-100, 0.5% sodium deoxycholate, 0.1% SDS, 50 mM Tris, pH 8.0; Thermo Fisher Scientific), supplemented with complete Protease Inhibitor Cocktail (Sigma Aldrich) (120 μL RIPA/1 × 10^6^ Cells). Protein concentrations were determined using Pierce BCA Protein Assay Kit (Thermo Fisher Scientific) as per manufacturer’s protocol. Ten μg of protein per sample was loaded on NovexTM 10-20% Tris-Glycine protein gels (Thermo Fisher Scientific) and transferred onto iBlot2 Transfer Stacks PVDF membrane (Thermo Fisher Scientific). Membranes were blocked with Odyssey Blocking buffer (Milennium Science, Victoria, Australia) for 2h, probed with rabbit anti-human LC3B (1:1000, Sigma L7543) and rabbit anti-human GAPDH (1:2500, Abcam 9485) overnight, followed by detection with goat anti-rabbit IgG DyLight 800 (1:20,000; Thermo Fisher Scientific). Bands were visualized using the Odyssey CLx system (LI-COR Biosciences, Nebraska, USA) and analyzed with Image Studio Lite V5.2 (LI-COR Biosciences).

### Immunofluorescence

Differentiated THP-1 cells were seeded onto a PDL coated, 96-well glass-bottom black tissue culture plate (4.5 × 10^4^ cells/well) and kept in RPMI-1640 medium minus phenol red (Thermo Fisher Scientific) supplemented with 10% heat-inactivated FBS at 37°C/5% CO_2_. Cells were infected with BCG, at a MOI of 10, with/without 100 nM 7α,25-OHC, with/without 10 μM GSK682753 for 2h, washed and incubated for a further 4h with agonists and antagonists. Cells were then fixed with 4% paraformaldehyde in PBS for 15 min, permeabilized with 0.05% saponin (Sigma Aldrich) for 20 min and blocked with 1% BSA, 0.05% saponin (Sigma Aldrich) for 1h. Cells were immunolabeled with rabbit anti-human LC3B (ThermoFisher L10382; 1:1000), 0.05% saponin at room temperature for 1h followed by Alexa FluorTM 647 goat anti-rabbit IgG (ThermoFisher A21245; 1:1000), 0.05% saponin at room temperature for 1h followed by nuclear staining with Hoechst 33342 (Thermo Fisher Scientific 62249; 1:2000) for 15 min. Cells were washed and confocal microscopy was performed using the Olympus FV3000, 60X magnification. Images obtained were analyzed with the ImageJ software [22].

### Murine GPR183 KO vs WT model

Equal numbers of male and female C57BL/6 WT and Gpr183^tm1Lex^ (age 18-20 weeks, 10 mice per group/timepoint) were aerosol infected with 300 CFU Mtb H_37_R_v_ using an inhalation exposure system (Glascol). At 2- and 5-weeks post infection, lungs and blood were collected for RNA and CFU determination. Formalin-fixed lung lobes were sectioned and examined microscopically and scored by a veterinary pathologist. Further details are available in the supplementary information.

### Statistical analysis

Statistical analysis was performed using GraphPad Prism v.7.0.3 (GraphPad Software). *T*-test and Wilcoxon’s test were used to analyze Nanostring data. Mann-Whitney *U* test and *t*-test were used to analyze in vitro infection, qPCR, and ELISA data. Data are presented as means ± SEM. Statistically significant differences between two groups are indicated in the figures as follows ns, *P* > 0.05; *, P < 0.05; **, P < 0.01; ***, P < 0.001; ****, P < 0.0001.

### Ethics statement

The human studies were approved by the Institutional Review Board of Stellenbosch University (N13/05/064 and N13/05/064A) and all study participants signed pre-approved informed consent documents prior to enrolment into the studies. All animal studies were approved by the Animal Ethics Committee of the University of Queensland (MRI-UQ/596/18) and conducted in accordance with the *Australian Code for the Care and Use of Animals for Scientific Purposes*.

## Results

### Blood GPR183 mRNA expression is reduced in patients with TB-T2D compared to TB patients without T2D

Blood was obtained from study participants with latent TB infection (LTBI, n=11), latent TB infection with T2D (LTBI+T2D, n=14), active pulmonary TB disease (TB, n=9), and active pulmonary TB disease with T2D (TB+T2D, n=7). Total RNA was extracted and NanoString analyzes performed. Among genes differentially expressed between TB and TB+T2D we identified a single GPCR, GPR183. We focused on GPR183 as GPCRs are *bona fide* drug targets due to their importance in human pathophysiology and their pharmacological tractability.

GPR183 expression was significantly down-regulated at diagnosis (p = 0.03, *t*-test) in blood from TB+T2D patients compared to TB patients without T2D (Figure 1A). The reduced GPR183 expression was not driven by diabetes *per se*, as there were no differences in GPR183 expression between LTBI and LTBI+T2D (Figure 1B). After 6 months, at the end of successful TB treatment, we saw GPR183 expression significantly increased (p=0.0156) in TB+T2D patients to a level comparable to the TB patients without T2D (Figure 1C). Therefore, we speculated that blood GPR183 expression is associated with extent of TB disease, which is frequently more severe in T2D patients. We indeed determined an inverse correlation between GPR183 mRNA expression in blood and TB disease severity on chest x-ray (Figure 1D).

**Fig 1.**
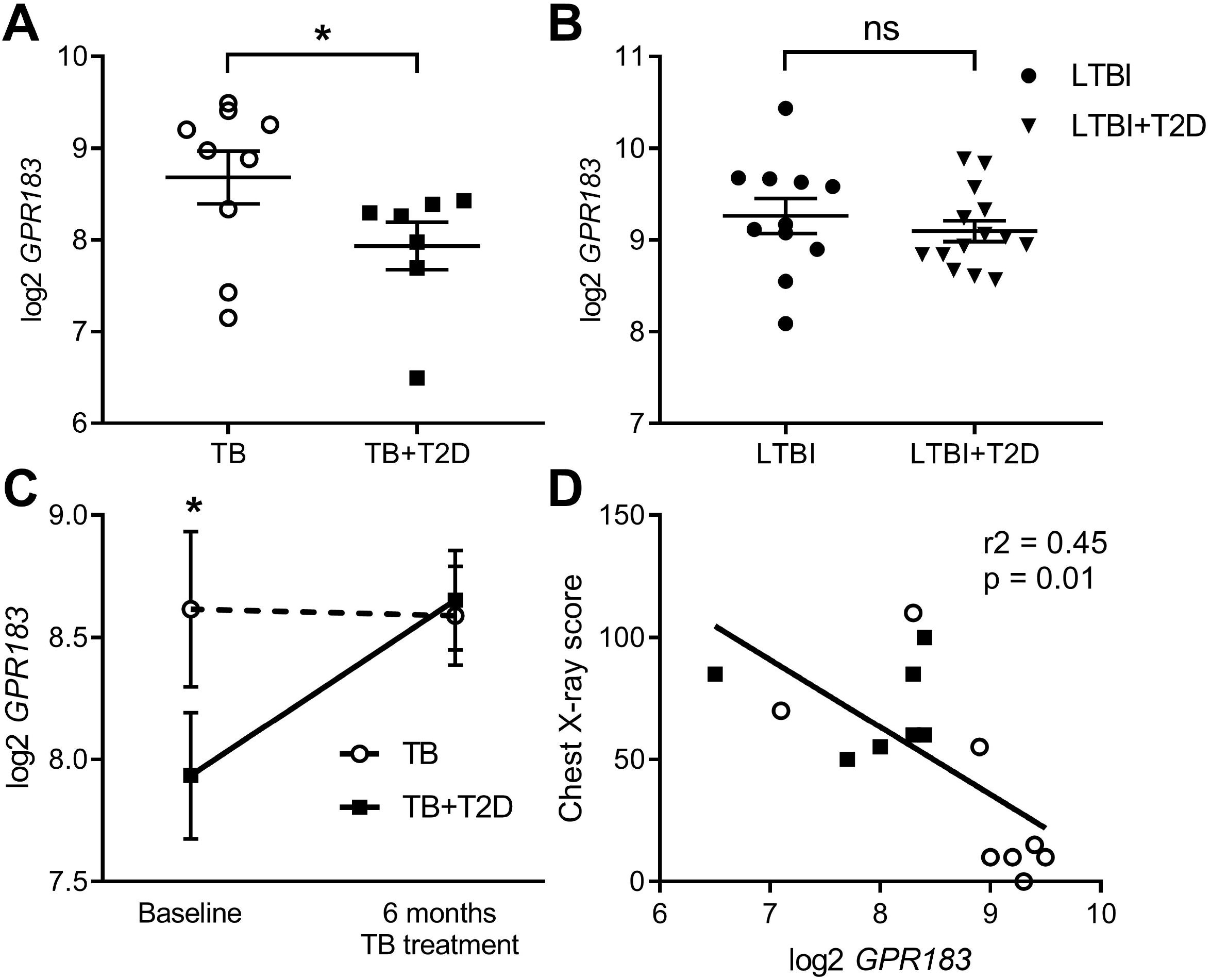
GPR183 mRNA expression in patients with active and latent TB infection with or without T2D. Total RNA was isolated from whole blood incubated overnight in QuantiFERON-TB Gold. *GPR183* mRNA expression was determined and normalized to reference genes using the NanoString technology. *GPR183* expression in whole blood of **(A)** TB (n=9) and TB+T2D (n=7) patients, **(B)** LTBI (n=11) and LTBI+T2D (n=14) patients, Wilcoxon test. **(C)** TB (n=9) and TB+T2D (n=7) patients at baseline and 6 month’s treatment, *t-test.* **(D)** Linear correlation between *GPR183* expression and chest × ray score, TB+T2D patients (n=7) filled squares, TB patients (n=8) open circles. Data are presented as means ± SEM; ns, *P* > 0.05; *, *P* ≤ 0.05.

In order to identify which cell type is associated with decreased expression of GPR183 in blood, we performed flow cytometry analysis for GPR183 expression on PBMCs from TB patients with and without T2D. We found that the only cell type with a significant reduction in GPR183 positivity in TB+T2D vs. TB, both in terms of frequency and median fluorescent intensity, was the non-classical monocyte population (Supplementary figure 1). We therefore next investigated whether GPR183 plays a role in the innate immune response during Mtb infection.

### Oxysterol-induced activation of GPR183 reduces intracellular mycobacterial growth

We investigated whether in vitro activation of GPR183 with its endogenous agonist impacts the immune response to mycobacteria in primary human MNs. MNs from 15 healthy donors were infected with BCG (n=7) or Mtb H_37_R_V_ (n=8) (Figure 2) at a MOI of 1 in the presence or absence of the GPR183 agonist 7α,25-OHC and/or the antagonist GSK682753. Activation of GPR183 by 7α,25-OHC significantly increased the uptake of BCG and Mtb H_37_R_V_ (Figure 2A) at 2h p.i. This increase in phagocytosis was abolished by the simultaneous addition of the GPR183 antagonist GSK682753, confirming that increased mycobacterial uptake was the result of GPR183 activation. Interestingly, we observed ~50% reduction in the growth of BCG and Mtb H_37_R_V_ (Figure 2B) by 48h p.i. in 7α,25-OHC treated cells, and again, this effect was abrogated by GSK682753. The addition of 7α,25-OHC and/or GSK682753 had no detrimental effect on the viability of human THP-1 cells (Supplementary figure 2A). There was also no effect of 7α,25-OHC and GSK682753 on BCG growth in liquid culture (Supplementary figure 2B), thus confirming that the significant mycobacterial growth inhibition in MN cultures was attributable to the immune modulatory activity of 7α,25-OHC via GPR183. Independently, we observed that H_37_R_v_ down-regulates GPR183 in primary MNs (Supplementary figure 3).

**Fig 2.**
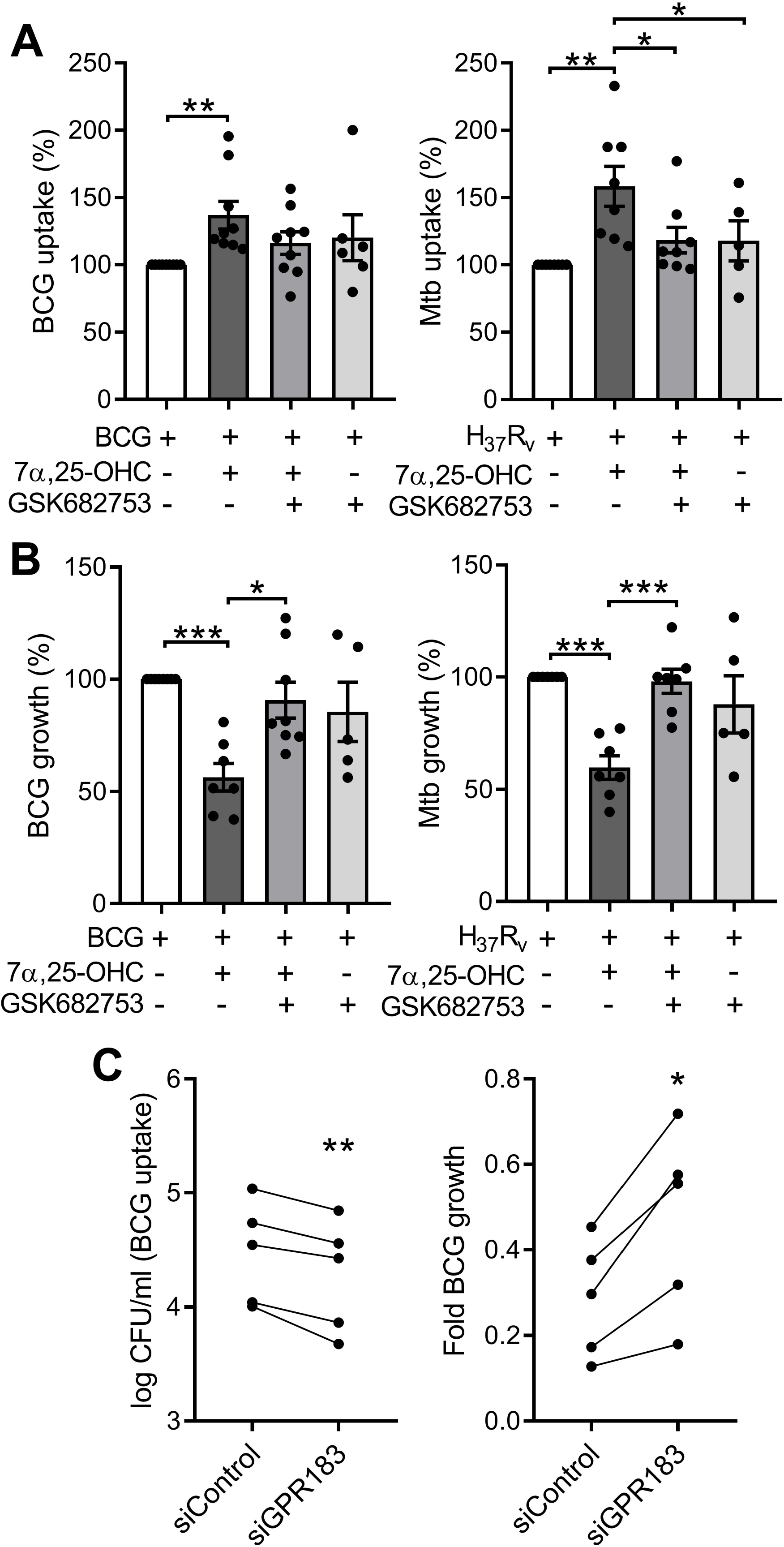
Oxysterol-induced activation of GPR183 in primary MNs significantly inhibits intracellular mycobacterial growth, while GPR183 knockdown increases intracellular mycobacterial growth. Primary MNs from eight donors **(A)** and seven donors **(B)** were infected with BCG or Mtb H_37_R_v_ (MOI 1), ± 7α,25-OHC (100 nM), ± GSK682753 (10 μM). Uptake of **(A)** BCG and Mtb H_37_R_v_ was determined at 2h p.i. Growth of **(B)** BCG and Mtb H_37_R_v_ was determined at 48h post-infection. Percent of mycobacterial growth was calculated as the fold change of CFU at 48h compared to CFU at 2h, normalized to non-treated cells. PMA-differentiated THP-1 cells were transfected with 20 nM of either negative control siRNA or GPR183 siRNA for 48h before infection with BCG (MOI 1). **(C)** Mycobacterial uptake was determined at 2h and **(D)** intracellular mycobacterial growth was determined at 48h p.i. (normalized to uptake). Data are presented as means ± SEM; *, *P* ≤ 0.05; **, *P* ≤ 0.01; ***, P ≤ 0.001; paired *t*-test.

To confirm the role of GPR183 in phagocytosis and growth inhibition, we next performed GPR183 siRNA knockdown experiments. Differentiated THP-1 cells were transfected with 20 nM of *GPR183*-targeting siRNA (siGPR183) or negative control siRNA (siControl). We observed ~80% reduction of *GPR183* mRNA level and ~50% reduction of protein expression in cells transfected with siGPR183 when compared to siControl-transfected cells (Supplementary figure 4A and B) at 48h. Forty-eight h after transfection the cells were infected with BCG at a MOI of 1. We observed a marked decrease in BCG uptake in cells transfected with siGPR183 (p = 0.0048) compared to siControl-transfected cells and a significant increase in intracellular mycobacterial growth over time (p = 0.0113, Figure 2C).

### GPR183 is a negative regulator of the type I interferon pathway in human MNs

In genome wide association studies GPR183 has been implicated as a negative regulator of the IRF7 driven inflammatory network [23]. Therefore, we focused subsequent experiments on type-I IFN regulation. To determine whether GPR183, a constitutively active GPCR [24], has a direct effect on *IRFs* and *IFNB1* expression we performed knockdown experiments in primary MNs. GPR183 knockdown (Supplementary figure 4C) up-regulated *IFNB1* (2.7-5.5 fold; *P* = 0.0115) as well as *IRF1, IRF3, IRF5* and *IRF7,* although the latter did not reach statistical significance (Figure 3A).

*IRF1*, *IRF5*, and *IRF7* transcripts were similarly up-regulated in whole blood from TB+T2D patients compared to TB patients (Figure 3B), consistent with the downregulation of *GPR183* mRNA expression (Figure 1C).

**Fig 3.**
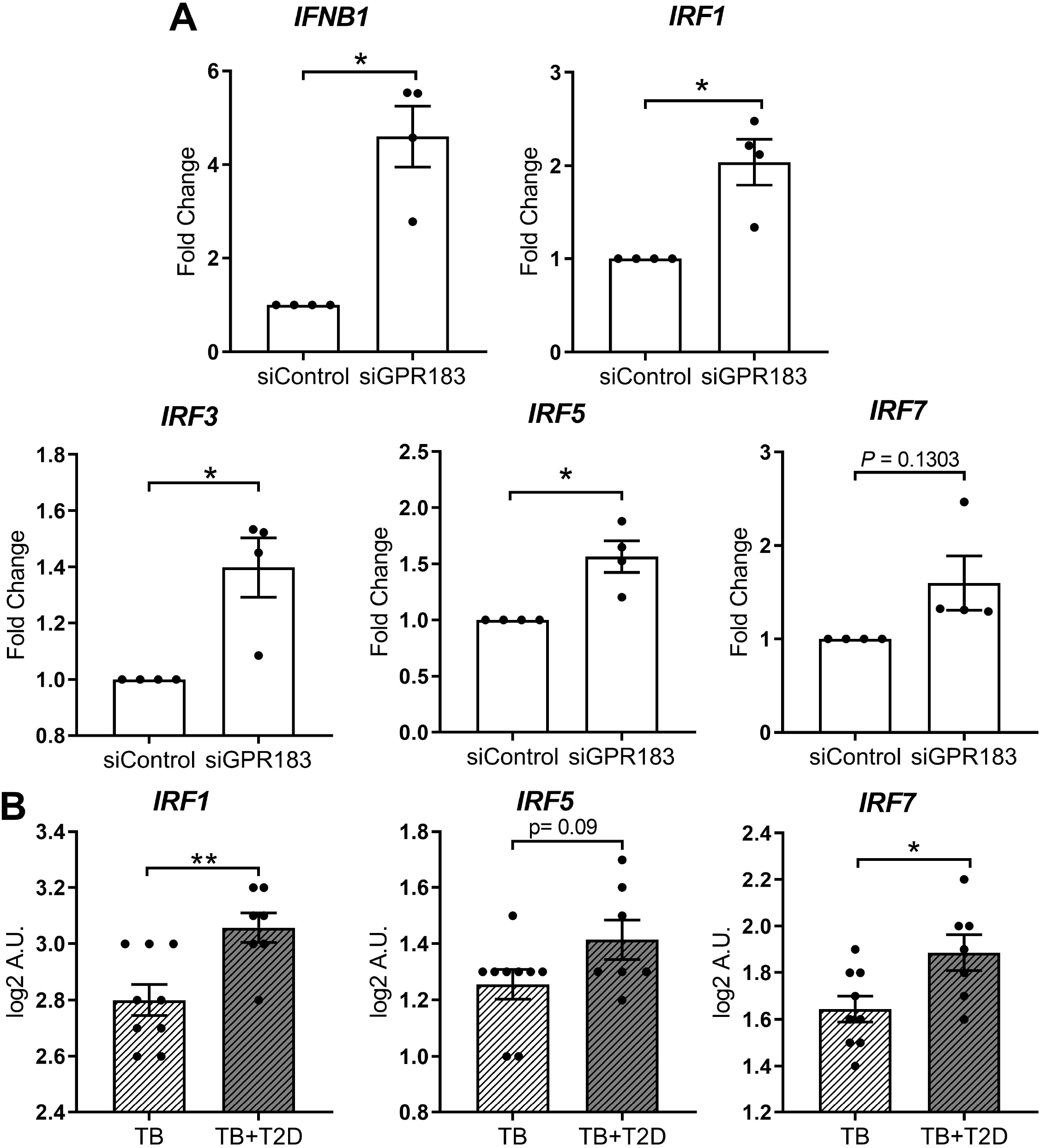
GPR183 knockdown increases expression of transcription factors regulating type I interferon responses. **(A)** Total RNA was isolated from primary MNs following 48h incubation with 20 nM GPR183 siRNA (or negative control siRNA). Gene expression of *IFNB1*, *IRF1*, *IRF3*, *IRF5, IRF7* was measured by qRT-PCR using RPS13 as reference gene. Data are, normalized to cells transfected with negative control siRNA. (**B**) NanoString analyses of RNA isolated from TB and TB+T2D cohort showed similar increase in type I IFNs associated genes *IRF1*, *IRF5*, *IRF7*. Data are presented as fold changes ± SEM; *, *P* ≤ 0.05; **, *P* ≤ 0.01; paired *t*-test.

### GPR183 activation induces a cytokine profile favoring Mtb control

Next, we investigated whether the reduced intracellular mycobacterial growth observed in primary MNs treated with 7α,25-OHC was associated with a change in MN secreted cytokines. Gene expression of *IFNB1, TNF*, and *IL-10* was measured 24h following infection with Mtb H_37_R_V_ at MOI of 1 (Figure 4A). The concentrations of the corresponding cytokines were measured in cell culture supernatant by ELISA (Figure 4B). Mtb infection significantly up-regulated the expression of *IFNB1* (*P* = 0.0068), *TNF* (*P* = 0.0001), *IL-10* (*P* < 0.0001) (Figure 4A) and *IL-1B* (Supplementary figure 5). 7α,25-OHC significantly down-regulated Mtb-induced *IFNB1* expression (*P* = 0.0017), while it did not affect *TNF*, *IL-10* or *IL-1B* expression. At the protein level, the concentrations of IFN-γ and IL-10, but not TNF-α or IL-1β were significantly lower in the culture supernatant of 7α,25-OHC-treated Mtb-infected primary MNs compared to untreated infected cells (*P* < 0.0001 and *P* = 0.0090, respectively, Figure 4B).

**Fig 4.**
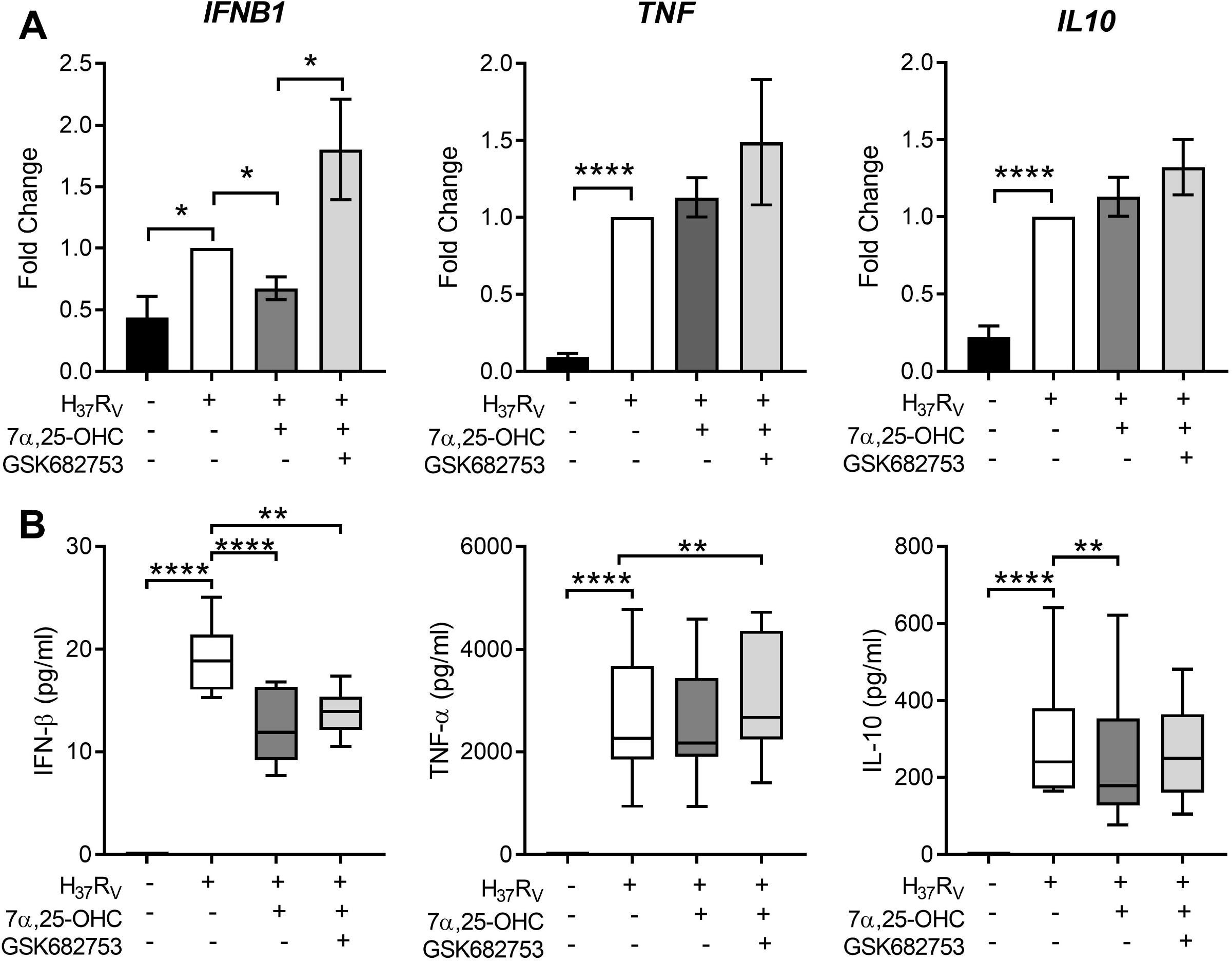
Activation of GPR183 leads to cytokine production favoring Mtb control. Primary MN from healthy donors (n=8) were infected for 2h with Mtb H_37_R_v_ (MOI 10:1), 7α,25-OHC (100 nM), and/or GSK682753 (10 μM). Cells were washed and left with drugs for a further 22h. Changes in the expression of **(A)** *IFNB1*, *TNF* and *IL10* were measured by qPCR and normalized to untreated infected cells. Concentrations of **(B)** IFN-β, TNF-α and IL-10 in the culture supernatant were measured by ELISA. Data are presented as mean fold change ± SEM or min to max for box plots; *, *P* ≤ 0.05; **, *P* ≤ 0.01; ****, *P* ≤ 0.0001; paired *t*-test.

### The oxysterol 7α,25-OHC induces autophagy

We aimed to identify whether 7α,25-OHC impacts the production of reactive oxygen species (ROS) and the autophagy pathway. ROS production in BCG-infected primary MNs was not affected by 7α,25-OHC (Supplementary figure 6); however, we observed an increase in accumulation of LC3B-II in BCG-infected THP-1 cells treated with 7α,25-OHC (*P* = 0.0119, Figure 5A). We next performed the experiments in absence and presence of the lysosomal inhibitor chloroquine in order to determine autophagic flux. Autophagic flux in BCG-infected cells was significantly increased with 7α,25-OHC treatment (*P* = 0.0069, Figure 5B). The simultaneous addition of the GPR183 antagonist GSK682753 with 7α,25-HC, decreased the levels of LC3B-II and autophagic flux, however, this did not reach statistical significance.

**Fig 5.**
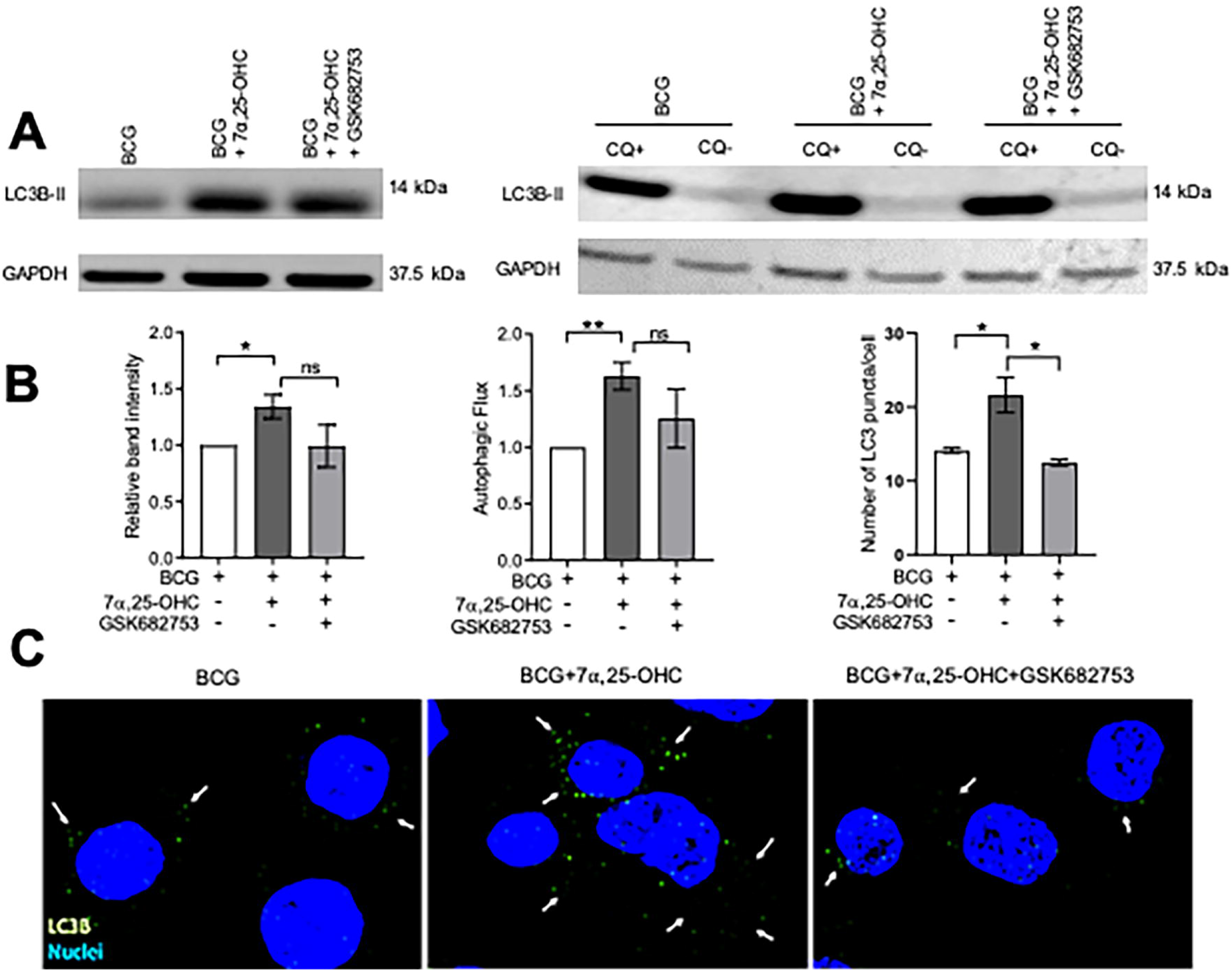
Treatment with 7 α,25-OHC induces autophagy. PMA-differentiated THP-1 cells were infected/uninfected and co-incubated with ± 7α,25-OHC, ± GSK682753, for 2h. Extracellular BCG was removed and cells were incubated for a further 4h or 22h in RPMI medium containing drugs. **(A)** Cells were lysed at 6h or 24h (Flux) p.i. **(B)** The band intensity was then normalized to the reference protein, GAPDH and further normalized to the BCG. Autophagic flux was obtained by subtracting chloroquine positive values with chloroquine negative values. **(C)** Cells were visualized using the Olympus FV 3000 confocal microscope. At least 30 cells were counted for every condition. Data are presented as ± SEM; ns, *P* > 0.05; *, *P* ≤ 0.05; **, *P* ≤ 0.01; unpaired *t*-test.

We next confirmed the induction of autophagy via microscopy. The number of LC3B-II puncta per cell increased in 7α,25-OHC stimulated BCG-infected THP-1 cells compared to untreated BCG-infected cells (*P* = 0.0358, Figure 5C). The 7α,25-OHC effect could be reduced by antagonist GSK682753 (*P* = 0.0196).

### GPR183 KO mice are unable to contain Mtb during the early stage of infection

To confirm the effect of the GPR183 receptor in vivo, we infected WT and GPR183 KO mice with aerosolized Mtb. At 2 weeks p.i., GPR183 KO mice showed significantly increased mycobacterial burden in the lungs compared to WT mice (*P* = 0.0084, Figure 6A), while the bacterial burden was comparable at 5 weeks p.i. (Supplementary figure 7). GPR183 KO mice also had higher lung pathology scores, although this did not reach significance (Figure 6B). GPR183 KO mice had significantly increased *Ifnb1* expression in the lungs (*P* = 0.0256; Figure 6C), along with increased *Irf3* (*P* = 0.0159), however, *Irf5* (Supplementary figure 8) and *Irf7* (Figure 6C) remained unchanged. *Irf7* transcription was increased in blood from GPR183 KO compared to WT mice (*P* = 0.0513; Fig 6D), but *Ifnb1*, *Irf3* and *Irf5* expression was not different (Figure 6D, Supplementary figure 6). At the RNA level *Tnf*, *Ifng* and *Il1b* were similar between GPR183 KO and WT mice (Figure 7A). Unexpectedly, at the protein level, the concentrations of IFN-β (*P* = 0.0232) and IFN-γ (*P* = 0.0232) were significantly lower in GPR183 KO mice lung, while TNF-α (*P* = 0.7394) and IL-1β (*P* = 0.0753) were similar to WT mice (Figure 7B).

**Fig 6.**
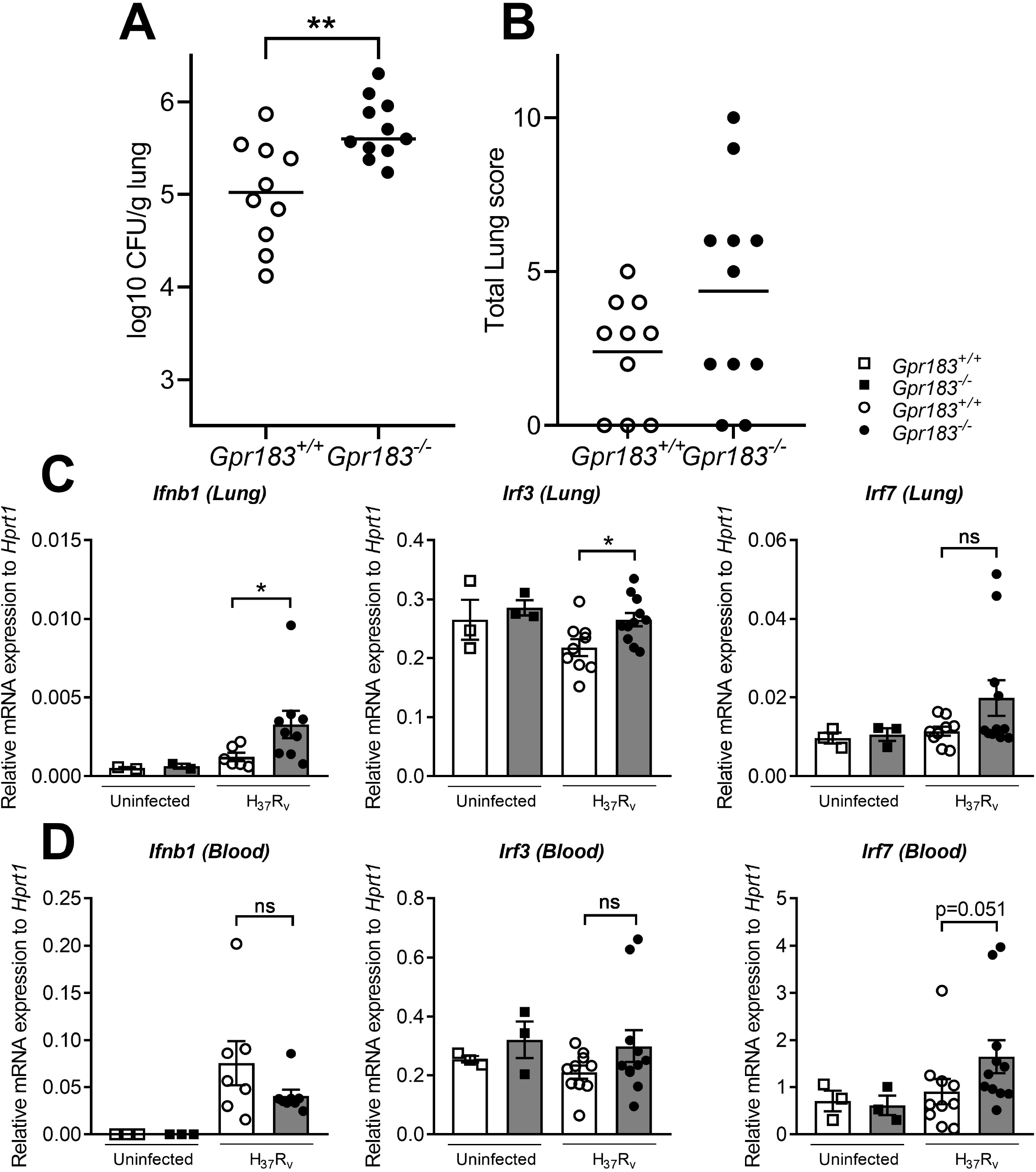
GPR183KO mice have higher lung CFU, corresponding with increased expression of transcription factors regulating type I interferon responses. Mice were infected with 300 CFU of aerosol Mtb H_37_R_v_. **(A)** Bacterial lung burden 2 weeks p.i. **(B)** Total histology lung score. RNA was isolated from Mtb-infected lung and blood samples 2 weeks p.i. **(C)** Gene expression of *Ifnb1*, *Irf3* and *Irf7* in the lungs, **(D)** *Ifnb1, Irf3* and *Irf7* in the blood, was measured by qRT-PCR using *Hprt1* as reference gene. Data are presented as ± SEM; ns, *P* > 0.05; *, *P* ≤ 0.05; **, *P* ≤ 0.01

**Fig 7.**
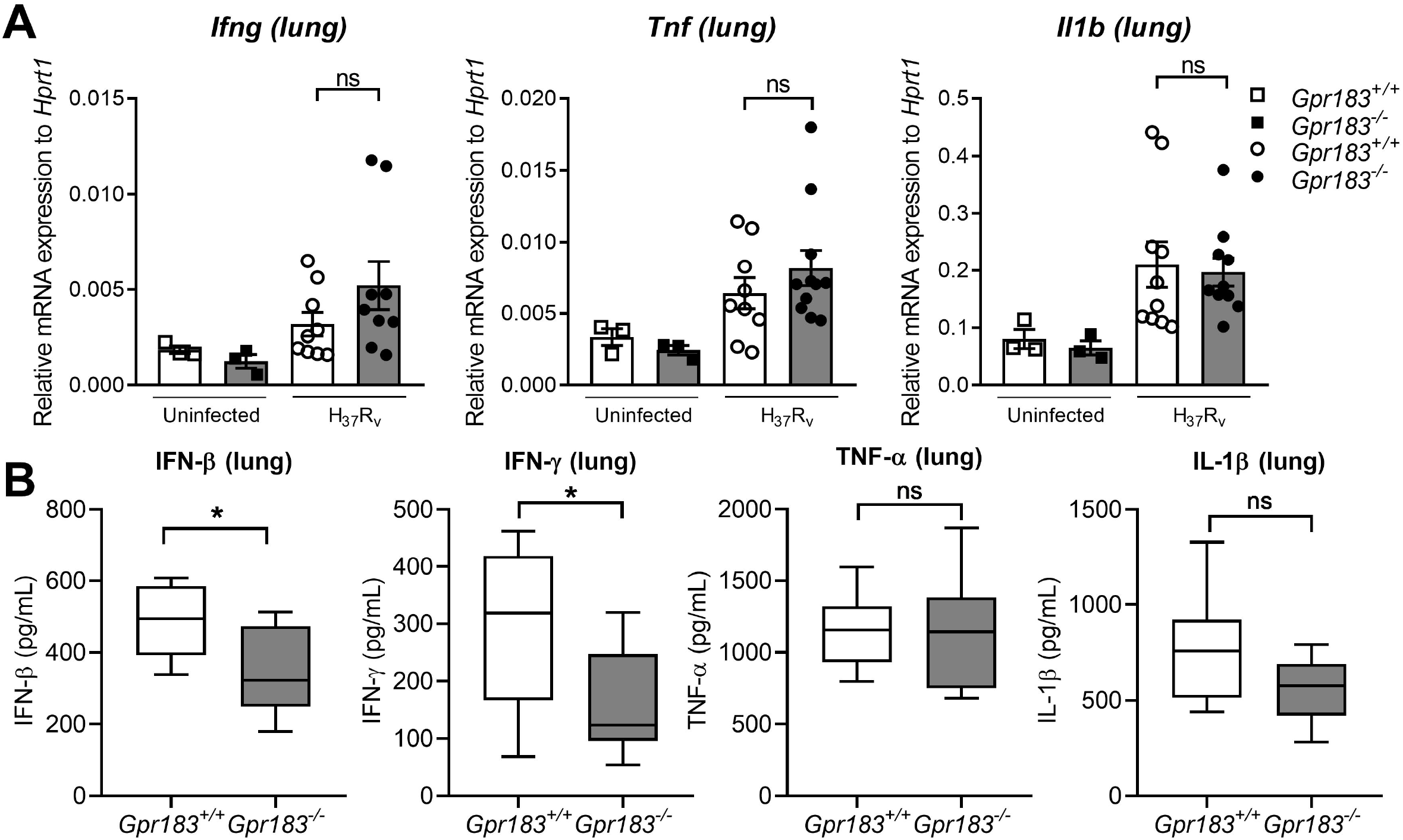
Pro-inflammatory cytokine expression at 2 weeks p.i. of Mtb H_37_R_v_-infected mice. Mice were infected with 300 CFU of aerosol Mtb H_37_R_v_. **(A)** Gene expression of *Ifng*, *Il1b* and *Tnf* in the lungs **(B)** Concentrations of IFN-β, IFN-γ, IL-1β and TNF-α in the culture supernatant were measured by ELISA. Data are presented as ± SEM; ns, *P* > 0.05; *, *P* ≤ 0.01

## Discussion

Historically oxidized cholesterols, so called oxysterols, were considered by-products that increase polarity of cholesterol to facilitate its elimination. However, they have recently emerged as important lipid mediators that control a range of physiological processes including metabolism, immunity, and steroid hormone synthesis [25].

Our findings define a novel role for GPR183 in regulating the host immune response during Mtb infection. We initially identified GPR183 through a blood transcriptomic screen in TB and TB+T2D patients and found an inverse correlation between GPR183 expression and TB disease severity on chest x-ray. Although we demonstrate that the decrease in blood GPR183 in TB+T2D patients is likely due, in part, to a decreased frequency of non-classical monocytes expressing GPR183, we cannot rule out that reduced GPR183 expression in whole blood is partially attributable to neutrophils and eosinophils, which are excluded from the PBMC population. In our study the TB patients with T2D had more severe TB compared to those without T2D, therefore we cannot ascertain whether lower GPR183 expression is linked to TB+T2D comorbidity or TB disease severity.

We demonstrate that activation of GPR183 by 7α,25-OHC in primary human MNs during Mtb infection results in significantly better control of intracellular Mtb growth. This is in contrast to a recently published study showing increased Mtb growth with 7α,25-OHC when added post-infection in murine RAW264.7 cells [26]. The discrepancies between the studies could also be attributed to the different cell types and infection dose, which was 25 times higher in the aforementioned study. Consistent with the findings of Tang et al. [26] in murine cells we show that mycobacterial infection down-regulates GPR183 in human MNs, which may be an immune-evasion strategy specific to mycobacteria since LPS, a constituent of Gram-negative bacteria, upregulates GPR183 [13]. Whether the observed increase in phagocytosis in the presence of 7α,25-OHC is a non-specific effect driven by internalization of agonist bound GPR183 and non-specific uptake of bacteria or an increase in pattern recognition receptors remains to be elucidated.

We further demonstrate that GPR183 activation by 7α,25-OHC reduces IFN-β expression and secretion in Mtb-infected primary MNs and targeted GPR183 knockdown significantly upregulating *IRFs* and *IFNB1*. Similarly, gene expression of *IRF1*, *IRF5*, and *IRF7* is up-regulated in TB+T2D patients compared to TB patients, and corresponds with down-regulation of *GPR183*, thereby demonstrating that GPR183 expression is associated with IFN regulatory factors during human TB and GPR183 is a negative regulator of type I IFNs in Mtb-infected human MNs.

There is mounting evidence that the production of type-I IFNs is detrimental during Mtb infection [27, 28]. Up-regulation of type-I IFN blood transcript signatures occur in TB disease and correlates with disease severity [29]. In macrophages, Mtb induces up-regulation of *IFNB1* expression as early as 4h p.i. to limit IL-1β production, a critical mediator in the host defense against Mtb [30]. Although 7α,25-OHC significantly reduced *IFNB1* mRNA, we did not observe an increase in *IL1B* mRNA, suggesting that the GPR183-mediated regulation of type-I IFN does not influence IL1B expression. In addition to GPR183 mediated reduction in IFN-β, we observed a decrease in IL-10 in Mtb-infected primary MNs treated with 7α,25-OHC. IL-10 production is induced by type-I IFN signaling [31, 32] and promotes Mtb growth [33] by reducing the bioavailability of TNF-α through the release of soluble TNF receptors and preventing the maturation of Mtb-containing phagosomes [33–36]. Collectively, we show that GPR183 is a negative regulator of type-I IFNs in primary MNs and agonist induced activation of GPR183 reduces Mtb-induced IFN-β production, while leaving expression of cytokines important for Mtb control unchanged.

Further confirming the role of GPR183, GPR183 KO mice infected with Mtb had significantly higher bacterial burden in the lung compared to WT mice 2 weeks p.i. (prior to initiation of the adaptive immune response to Mtb) with this effect disappearing at 5 weeks p.i., when T cell responses against Mtb are fully established. Our results thus strengthen the contention that GPR183 plays an important role in the innate immune control of Mtb irrespective of hyperglycemia. We confirmed the importance GPR183 in regulating type-I interferons during Mtb infection in vivo. GPR183 KO mice infected with Mtb had significantly increased lung *Ifnb1* and *Irf3* mRNA. Unexpectedly, IFN-β and IFN-γ secretion were both significantly downregulated in the lung. These differences between mRNA and protein levels may be due to kinetic parameters of transcription versus translation or mRNA stability versus protein consumption.

Furthermore, we demonstrate that the GPR183 agonist 7α,25-OHC promotes autophagy in macrophages infected with mycobacteria. Autophagy is a cellular process facilitating the elimination of intracellular pathogens including Mtb [37]. Antimicrobial autophagy was shown to be inhibited by *Mycobacterium leprae* through upregulation of IFN-β and autocrine IFNAR activation which in turn increased expression of the autophagy blocker OASL (2’-5’-oligoadenylate synthetase like) [38]. Whether there is a link between the 7α,25-OHC-induced reduction of IFN-β production and the increase in autophagy remains to be investigated in future studies.

Several autophagy promoting re-purposed drugs including metformin are currently being assessed as HDTs for TB [39]. We propose that GPR183 is a potential target for TB HDT, warranting the development of specific, metabolically stable small-molecule agonists for this receptor to ultimately improve TB treatment outcomes in TB patients with and without T2D co-morbidity.

## Author contributions

ATG, SB and KR wrote the manuscript; ATG, SB, RS, SH, HS, MDN, CXF, LK, HT, TW, HBO, AMH, CEM, LVVC, NPW carried out the experiments; ATG, SB, MD, HS, RS and SH analyzed the data; TMP, MMR, LSS, GW, KR interpreted the data and developed the theoretical framework, KR conceived the original idea; all authors provided critical feedback and helped shape the research, analysis and manuscript.

## Acknowledgements

We thank the clinical research team at Stellenbosch University for assistance with identification and recruitment of study participants as well as coordination of clinical and administrative activities. We thank Matthew Sweet for critical review of the manuscript. Illustrations were created with Biorender.com.

