## Supplemental data for "GPR183 regulates interferons and bacterial growth during *Mycobacterium tuberculosis* infection: interaction with type 2 diabetes and TB disease severity"

**Methods**

*Study participant recruitment, diagnosis and treatment.*

Briefly, the percentage of affected lung was graded using visual estimation of the extent of opacification, cavitation or pathology by CXR, with dense consolidation demonstrating a percentage of 90 – 100% for a quadrant, while less dense infiltrates attracting a lesser percentage. The total score of the percentage of affected lung (%) + the cavitation score (40 if present or 0 if not) was used to determine the overall chest x-ray score. All TB patients were screened for T2D at baseline and month 6 by laboratory-based HbA1c assay and, individuals with an HbA1c ≥ 6.5% and random plasma glucose ≥ 200 mg/dL at both time points were considered to have TB-T2D co-morbidity. Baseline samples were collected prior to treatment. All TB patients had drug-susceptible pulmonary TB and were on standard TB treatment (isoniazid, rifampicin, ethambutol and pyrazinamide for the first two months, followed by isoniazid and rifampicin for four months) in accordance with the South African National Tuberculosis Program. All TB patients irrespective of T2D co-morbidity completed > 80% of their TB medication and were cured according to WHO criteria at the end of treatment.

Participants with LTBI were close contacts of TB patients, recruited as part of the NIH funded ALERT study and tested positive on QuantiFERON-TB Gold in tube assay. T2D status was determined as described above for TB patients. Participants were excluded from the study if they were pregnant, HIV positive, had drug-resistant Mtb or were using immunosuppressive drugs.

*Quantiferon-TB gold in tube (QFT) assay*

One mL of blood, collected in a LiHep tube (1x 4 mL), was transferred to each of the three QFT tubes (nil tube containing no additives; Mtb-Ag tube coated with an antigen cocktail including ESAT 6, CFP-10 and TB 7.7 specific for Mtb and, the mitogen tube containing phytohaemagglutin-P (PHA)) of the QFT assay (Qiagen, Hilden, Germany) and processed following the manufacturer’s instructions. After incubation at 37°C for 16-22h, blood was transferred from the QFT tubes into 2 mL cryogenic vials (Sigma-Aldrich, Missouri, USA) and centrifuged for 15 min at 15,000 x g. The supernatants were stored at -80°C and the pellet stored in RNA*Later* (Thermo Fisher Scientific, Virginia, USA) at 4°C overnight, before being transferred to -80°C. The preserved QFT pellets were used for RNA extractions and the supernatants used to measure IFN-γ concentrations by ELISA.

*siRNA transfection*

Delivery of GPR183 siRNA (Silencer Select siRNA #s4430, Thermo Fisher Scientific) into primary MN or differentiated THP-1 cells was performed using Lipofectamine RNAiMAX (Thermo Fisher Scientific) in a 96-well format with 1.3 x 10^5^ cells/well or 6 x 10^4^ cells/well, respectively. Cells were incubated with 0.6 μL Lipofectamine RNAiMAX and 20 nM GPR183 siRNA or Silencer Select Negative Control No. 1 siRNA (Thermo Fisher Scientific) in OptiMEM I Reduced Serum medium (Thermo Fisher Scientific) for 48h before further experiments. Knockdown efficiency was measured by qPCR and by flow cytometry 48h after transfection.

*RNA Isolation, cDNA synthesis and Quantification of gene expression*

MNs infected with Mtb H_37_R_v_ were lysed by adding 500 μl of TRIzol reagent. Subsequently, 100 μl of cold chloroform was added, followed by centrifugation at 12,000 x *g*, 4^o^C, and the resulting upper aqueous RNA-containing phase was collected for further RNA isolation. Total RNA was isolated using ISOLATE II RNA Mini kit (Bioline, London, UK) according to the manufacturer’s protocol with on-column DNase digestion. RNA quantity and the 260/280 and 260/230 ratios were measured using nanodrop spectrophotometer. Up to 1 μg of RNA from each sample was reverse transcribed using Tetro cDNA Synthesis Kit (Bioline) according to manufacturer’s protocol (45^o^C for 30 min, 85^o^C for 5 min). The cDNA product was stored at -20^o^C until further use. Quantification of gene expressions was performed with the QuantStudio 7 Flex Real-Time PCR system (Thermo Fisher Scientific) using SensiFAST SYBR Lo-ROX kit (Bioline). A list of primers used can be found in S1 Table.

Cycle conditions were 95^o^C for 10 min, followed by 40 cycles of 95^o^C for 15 s and 60^o^C for 30 s. Melt curves were obtained by gradually increasing the temperature from 60^o^C to 95^o^C with 0.05^o^C/s increment.

*Flow cytometry*

THP-1 cells were detached using PBS containing 5 mM EDTA 48h after siRNA transfection and incubated with human FcR binding reagent (eBioscience) for 15 min. Mouse anti-human GPR183 BV421 antibody (Biolegend, 368910) was added and cells were incubated on ice for an additional 20 min. Cells were washed and resuspended in FACS buffer (PBS, 1% FBS, 2 mM EDTA) and events acquired on a LST Fortessa X-20 flow cytometer (BD Biosciences). Data were analyzed using FACS DIVA.

*Cytokine measurements*

Concentrations of cytokines (TNF-α, IL-1β, IFN-γ, IFN-β, IL-10) secreted into the culture supernatant in response to Mtb H_37_R_v_ infection and cell free supernatants from lung homogenates of *Mtb*-infected mice were measured by DuoSet ELISA Kit (R&D Systems) according to the manufacturer’s protocol.

*Histopathology*

The following features were assessed individually and combined for a total lung score: peribronchiolitis, perivascular leukocyte infiltration (“perivasculitis”), alveolitis, “granuloma” formation (i.e., granulomatous inflammation), and necrosis, as described in details previously [1], using a Nikon Eclipse 50i microscope with objectives ranging from 2x-60x.

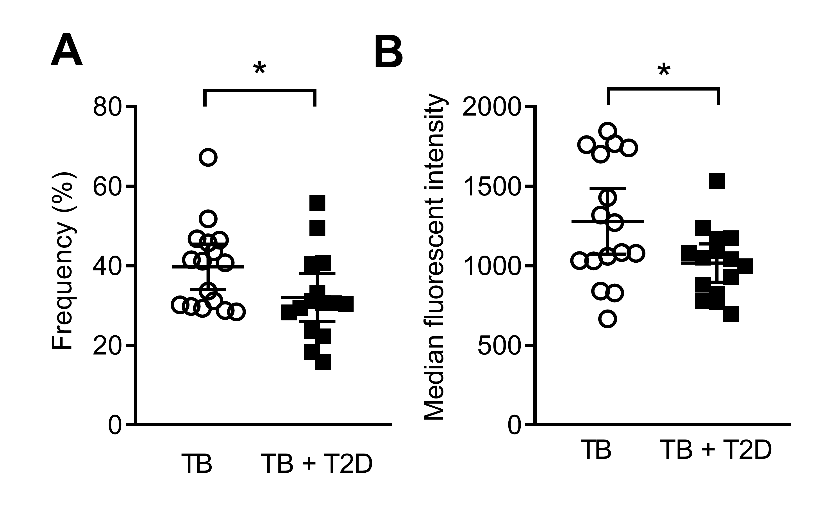

**Supplementary figure 1. Expression of GPR183+ non classical monocytes in blood from TB and TB-T2D patients.** Flow cytometry analysis was performed on PBMCs of participants demonstrating a down regulation of GPR183 expression in **(A)** frequency **(B)** median fluorescent intensity on non-classical monocytes (GPR183^+^ CD14^-^ CD16^+^) in TB+T2D patients compared to TB patients. Data are presented as ± SEM; *, *P* ≤ 0.05.

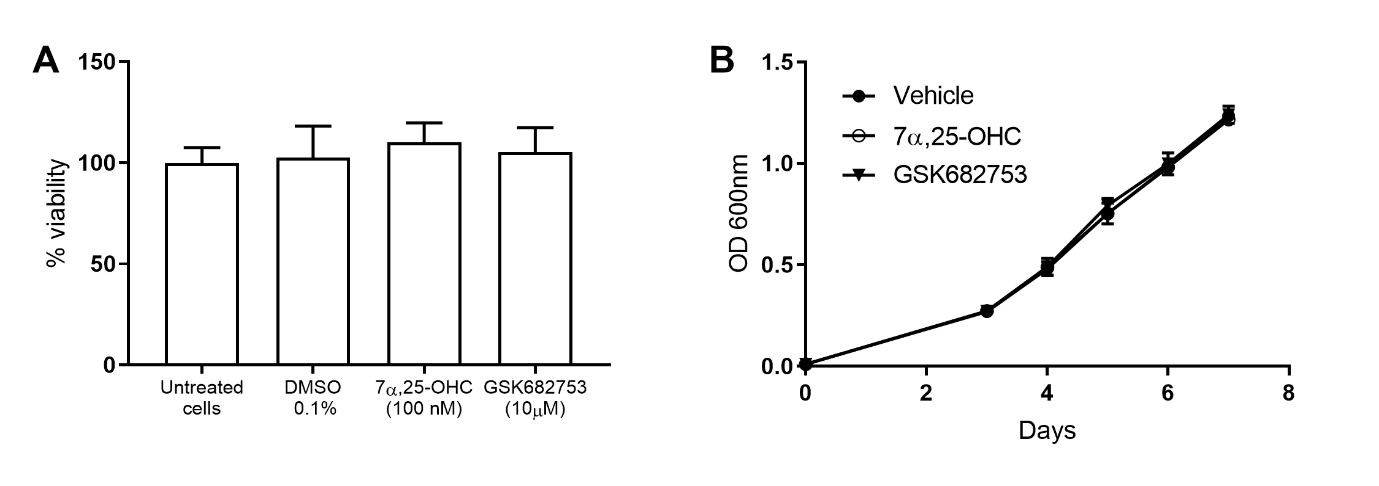

**Supplementary figure 2. Co-incubation with** **7α,25-OHC or GSK682753 does not affect the viability of MNs or Middlebrook 7H9 broth culture growth of BCG.** **(A)** PMA-differentiated THP-1 were co-incubated with 7α,25-OHC (100 nM) or GSK682753 (10 μM) for 24h before cell viability was measured by adding Alamar Blue reagent. No significant changes in cell viability were observed (p>0.05, Mann-Whitney U test). **(B)** 7 day growth curve of BCG co-incubated with 7α,25-OHC (100 nM) or GSK682753 (10 μM) or vehicle (0.1% DMSO) in Middlebrook 7H9 broth culture (7H9, 10% Oleic Albumin Dextrose Catalase, 0.02% glycerol, 0.05% Tween 80). BCG was adjusted to an OD600 of 0.01 prior to the addition of the compounds. Cell density was measured daily in triplicate. No significant changes in cell viability were observed (*P* > 0.05, Mann-Whitney U test). Data are presented as means ± SEM.

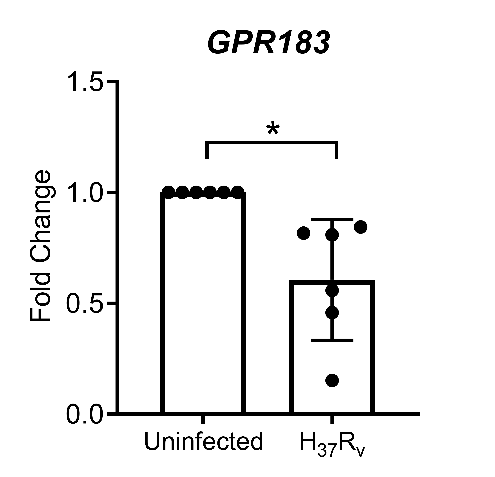

**Supplementary figure 3.** Primary human MNs from healthy donors (n=16) were incubated for 2h with live Mtb H_37_R_v_ (MOI:1), washed and left for a further 22h. GPR183 expression is down-regulated in mRNA. Data are presented as means ± SEM of six independent replicates; *, *P* ≤ 0.05.

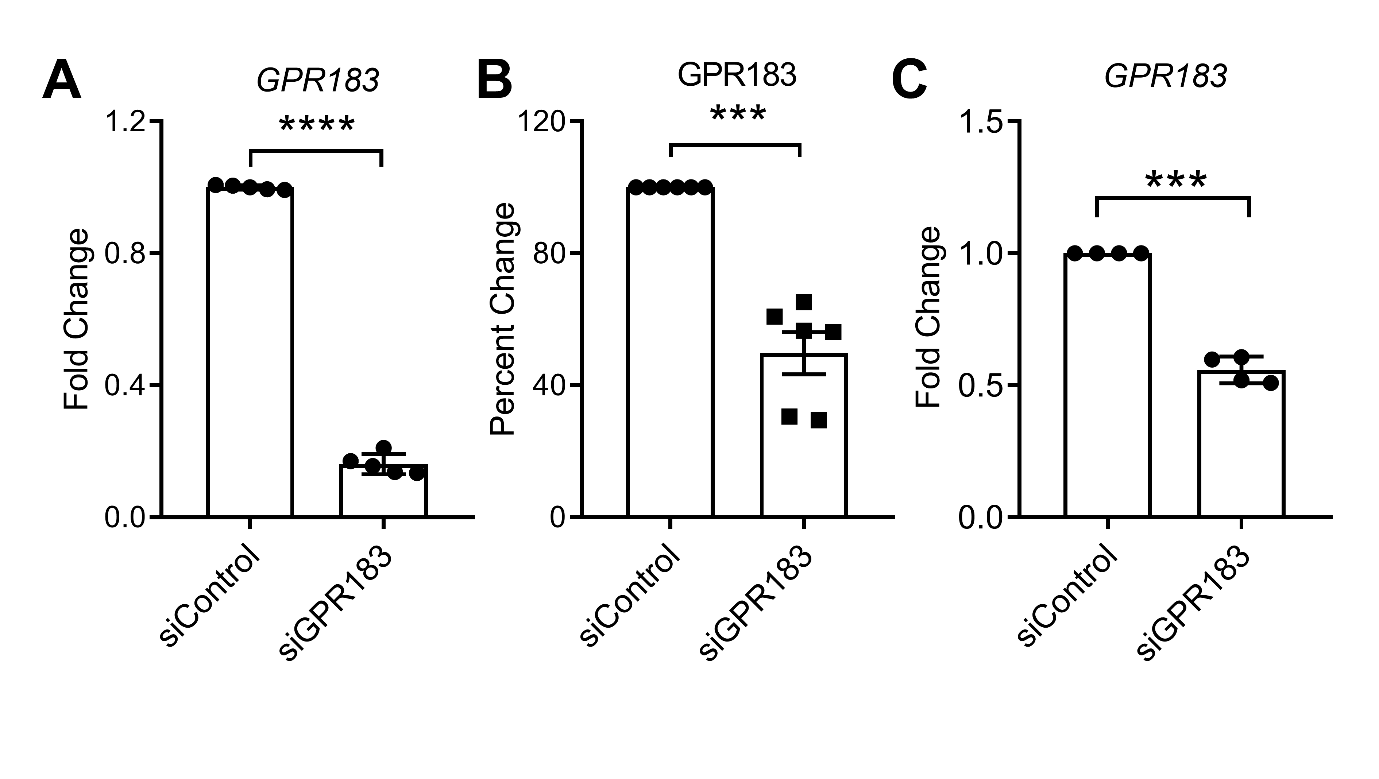

**Supplementary figure 4. (A)** PMA-differentiated THP-1 cells and primary MNs were transfected with 20 nM of either negative control siRNA or GPR183 siRNA for 48h before infection with BCG at MOI 1. Efficiency of the GPR183 knockdown was determined at **(A)** THP-1 cells RNA level by qPCR and **(B)** THP-1 cells protein level by flow cytometry, **(C)** Primary MNs RNA levels by qPCR. Data are presented as means ± SEM; ***, P ≤ 0.001; ****, *P* ≤ 0.0001

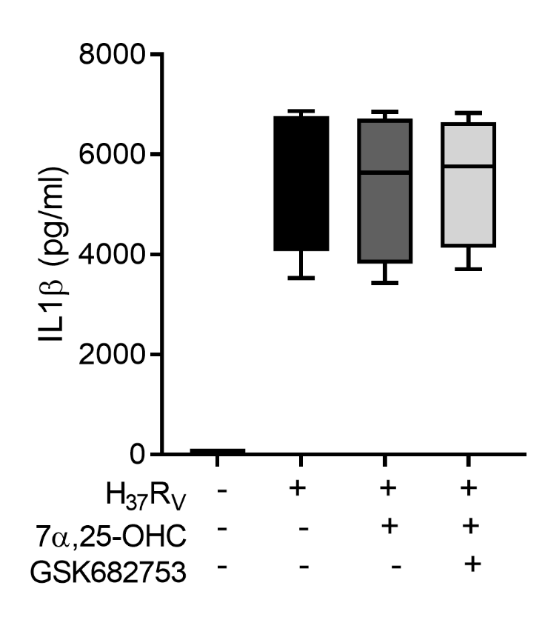

**Supplementary figure 5. Activation of GPR183 leads to cytokine production.** Primary MN from healthy donors (n=8) were infected for 2h with Mtb H_37_R_v_ (MOI 10:1), 7α,25-OHC (100 nM), and/or GSK682753 (10 μM). Cells were washed and left with drugs for a further 22h. Concentrations of IL-1β in the culture supernatant were measured by ELISA. Data are presented as mean fold change ± SEM or min to max for box plots; ns, *P* > 0.05.

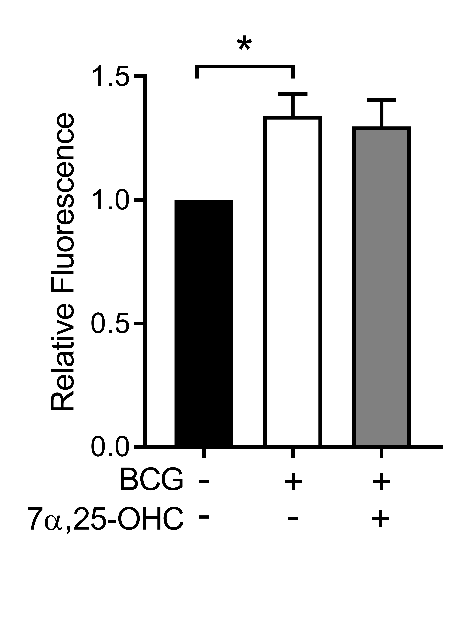

**Supplementary figure 6.** **Treatment with 7α,25-OHC does not affect Reactive Oxygen Species (ROS) production in primary MNs.** Primary MNs (n=3) were infected with BCG (MOI 10) and co-incubated with 7α,25-OHC for 2h. Cells were then washed twice to remove non-phagocytosed BCG, and cells were stained with H_2_DCFDA for 1h. Following this, DCFDA was replaced with RPMI containing 7α,25-OHC and cells were further incubated for 40 min before fluorescence intensity was read. Readings were corrected to OD600 values, blank corrected, and normalised to unstimulated. Data are presented as ± SEM; ns *P* > 0.05; *, *P* ≤ 0.05.

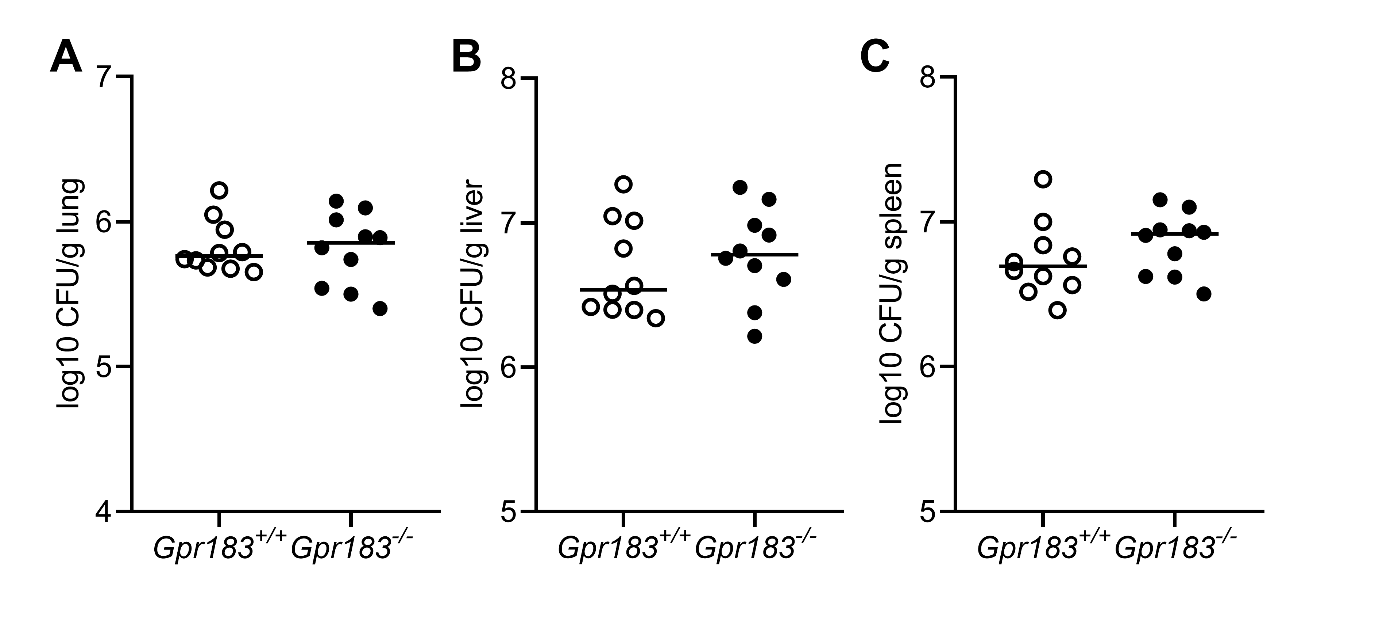
 **Supplementary figure 7.** Mice were infected with 300 CFU of aerosol Mtb H_37_R_v_. Bacterial burden was examined 5 weeks p.i **(A)** Lung burden **(B)** Liver burden **(C)** Spleen burden. Data are presented as ± Mean; ns, *P* > 0.05

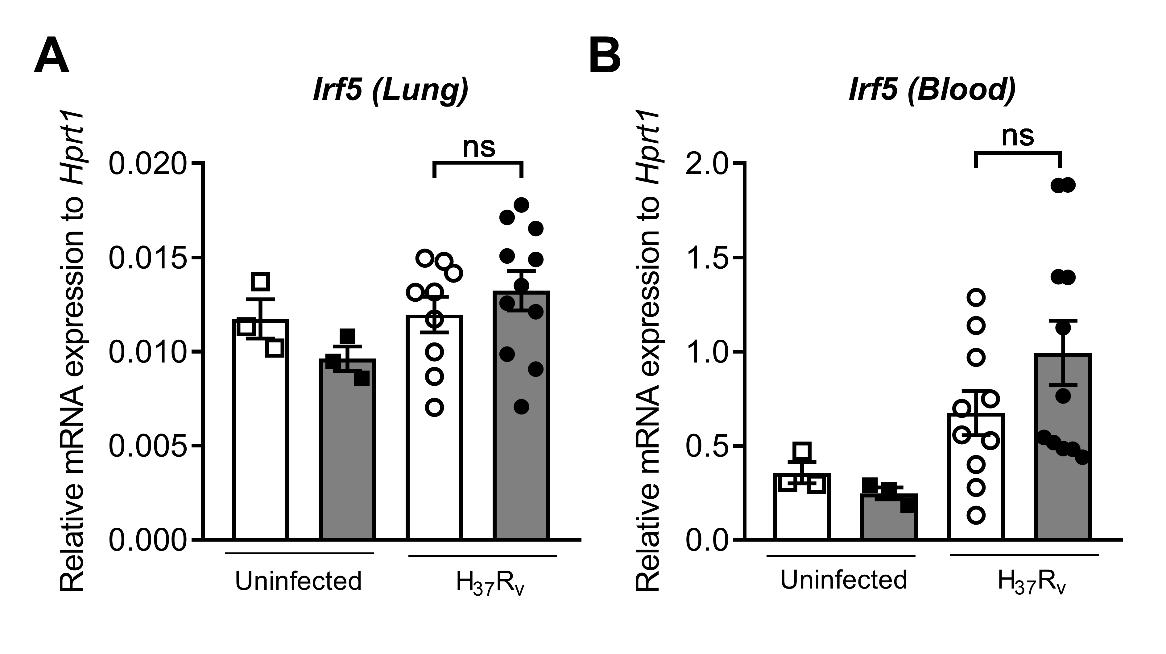

**Supplementary figure 8.** **Gene expression of Mtb H_37_R_v_-infected mice.** Mice were infected with 300 CFU of aerosol Mtb H_37_R_v_. RNA was isolated from Mtb-infected lung and blood samples 2 weeks p.i. Gene expression of *Irf5* in the lungs and blood measured by qRT-PCR using *Hprt1* as a reference gene. Data are presented as mean fold change ± SEM or min to max for box plots. Data are presented as ± SEM; ns, *P* > 0.05.

**Supplementary Table 1.**

**List of primers**:

Human GPR183

forward primer 5’-ACCACCGCTTTGCCTACAC GAA-3’

reverse primer 5’-CAC CACAGCAATGAAGCGGTCA-3’;

Human TNF

forward primer 5’-CTCTTCTGCCTGCTGCACTTTG-3’

reverse primer 5’-ATGGGC TACAGGCTTGTCACTC-3’;

Human IL10

forward primer 5’-TCTC CGAGATGCCTTCAG CAGA-3’

reverse primer 5’-TCAGACAAGGCTTGGCAACCCA-3’;

Human IFNB1

forward primer 5’-ACGCCGCATTGACCATCTAT-3’

reverse primer 5’-GTCTCATTCCAGCCAGTGCT-3’;

Human IRF1

forward primer 5’-GAGGAGGTGAAAGACCAGAGCA-3’

reverse primer 5’-TAGCATCTCGGCTGGACTTCGA-3’;

Human IRF3

forward primer 5’-TCTGCCCTCAACCGCAAAGAAG-3’

reverse primer 5’-TACTGCCTCCACCATTGGTGTC-3’;

Human IRF5

forward primer 5’-TATGCCATCCGCCTGT GTCAGT-3’

reverse primer 5’-GCCCTTTTGGAACAGGATGAGC-3’;

Human IRF7

forward primer 5’-CCACGCTATACCATCTACCTGG-3’

reverse primer 5’-GCTGCTATCCAGGGAAGACACA-3’;

Human RPS13

forward primer 5’-GCCTT ACTCCTTCACAGATCGG-3’

reverse primer 5’-GGAAGATCAGGAGCA AGTCCCT-3’;

Mouse *Hprt*

forward primer 5’-CCCCAAAATGGTTAAGGTTGC-3’

reverse primer 5’- AACAAAGTCTGGCCTGTATCC-3’;

Mouse *Ifnb1*

forward primer 5’- GTCCTCAACTGCTCTCCACT-3ʹ

reverse primer 5’- CCT GCAACCACCACTCATTC-3ʹ.

Mouse *Ifng*

forward primer 5’- CAGCAACAGCAAGGCGAAAAAG-3’

reverse primer 5’- TTTCCGCTTCCTGAGGCTGGAT-3’

Mouse *Irf3*

forward primer 5’- CGGAAAGAAGTGTTGCGGTTAGC-3’

reverse primer 5’- CAGGCTGCTTTTGCCATTGGTG-3’

Mouse *Irf5*

forward primer 5’- CCTACAGAACCACTCTTGCCTG-3’

reverse primer 5’- CCTTGTGGGTTGCTAGTGGTGA-3’

Mouse *Irf7*

forward primer 5’- CCTCTGCTTTCTAGTGATGCCG-3’

reverse primer 5’- CGTAAACACGGTCTTGCTCCTG-3’

Mouse *Tnfa*

forward primer 5’-TAGCCCACGTCGTAGCAAAC-3’

reverse primer 5’- ACAAGGTACAACCCATCGGC-3’

Mouse *Il1b*

forward primer 5’- TGGACCTTCCAGGATGAGGACA-3’

reverse primer 5’- GTTCATCTCGGAGCCTGTAGTG-3’

1. Flores-Valdez MA, Pedroza-Roldan C, Aceves-Sanchez MJ, et al. The BCGDeltaBCG1419c Vaccine Candidate Reduces Lung Pathology, IL-6, TNF-alpha, and IL-10 During Chronic TB Infection. Front Microbiol **2018**; 9:1281.
